# Tumor establishment requires tumor autonomous and non-autonomous deregulation of homeostatic feedback control

**DOI:** 10.1101/541912

**Authors:** Sang Ngo, Jackson Liang, Yu-Han Su, Lucy Erin O’Brien

## Abstract

**Summary:** In healthy adult organs, robust feedback mechanisms control cell turnover to enforce homeostatic equilibrium between cell division and death [1, 2]. Nascent tumors must subvert these mechanisms to achieve cancerous overgrowth [3–7]. Elucidating the nature of this subversion can reveal how cancers become established and may suggest strategies to prevent tumor progression. In adult *Drosophila* intestine, a well-studied model of homeostatic cell turnover, the linchpin of cell equilibrium is feedback control of the EGF protease Rhomboid (Rho). Expression of Rho in apoptotic cells enables them to secrete EGFs, which stimulate nearby stem cells to undergo replacement divisions [8]. As in mammals, loss of *adenomatous polyposis coli* (APC) causes *Drosophila* intestinal stem cells to form adenomas [9]. Here we demonstrate that *Drosophila APC*^−/−^ tumors trigger widespread Rho expression in non-apoptotic cells, resulting in chronic EGF signaling. Initially, nascent *APC*^−/−^ tumors induce *rho* in neighbor wild-type cells via acute, non-autonomous activation of JNK. During later growth and multilayering, *APC*^−/−^ tumors induce *rho* in tumor cells by autonomous downregulation of E-cadherin (E-cad) and consequent activity of p120-catenin. This sequential dysregulation of tumor non-autonomous and -autonomous EGF signaling converts tissue-level feedback into feed-forward activation that drives cancerous overgrowth. Since Rho, EGFR, and E-cad are associated with colorectal cancer in humans [10–17], our findings may shed light on how human colorectal tumors progress.

## Results

To investigate how tumors subvert cell equilibrium in the *Drosophila* intestine (midgut) (Figure S1A), we used low-frequency, *hs-flp*-mediated MARCM recombination [18] to generate stem cells that were (1) marked by heritable fluorescent protein expression and (2) either control genotype or homozygous for null alleles of *Drosophila Apc1* and *Apc2* (hereafter, *APC^−/−^*) [19, 20] (Figure S2A). We allowed these marked stem cells to form multicellular clones and examined clone size and morphology (*c.f.* STAR Methods, “Clone visualization and quantification”). Midguts from mated females were used exclusively here and in subsequent experiments.

As described previously [21–25], *APC*^−/−^ stem cells frequently gave rise to large, multilayered adenomas over time (Figures 1A, S1B-F). Whereas most 21-day control clones contained fewer than 50 cells, many 21-day *APC*^−/−^ clones contained 100-500 cells (Figure 1G). This tumorous growth was accompanied by epithelial multilayering. At two days after induction, nearly all *APC*^−/−^ clones were single-layered (Figure S1F); by 21 days after induction, in contrast, 36.2 ± 1.7% of *APC*^−/−^ clones were multilayered. These overgrown, multilayered *APC*^−/−^ masses protruded conspicuously into the midgut lumen (Figures S1C, S1D, S1E, S1G), reminiscent of *APC-*inactivated colonic adenomas in humans [26].

**Figure 1.**
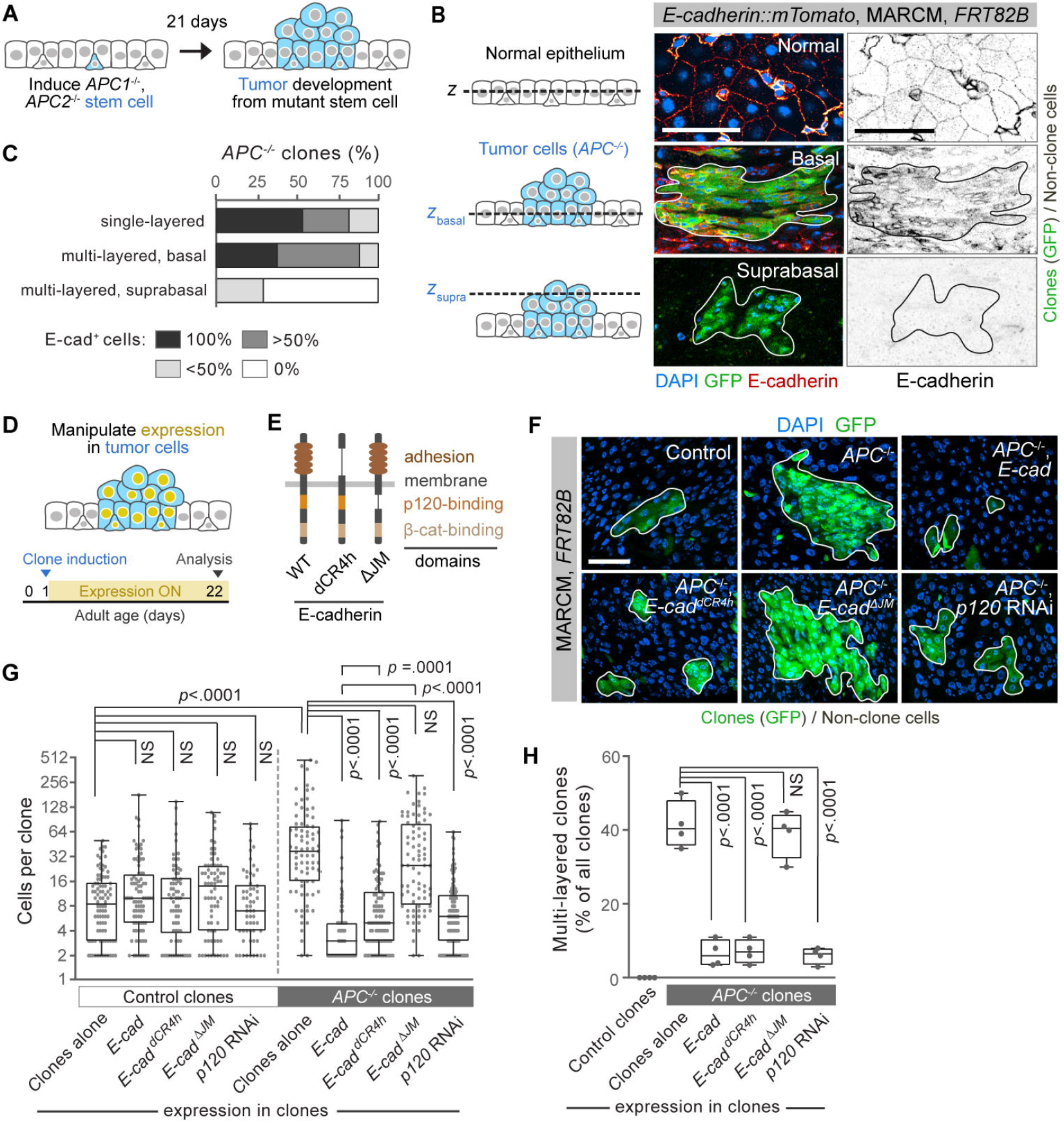
Growth of multilayered *APC*^−/−^ tumors requires tumor-autonomous downregulation of E-cadherin (E-cad) and consequent deregulation of p120-catenin. **(A)** Experimental schema for B and C. Sparsely distributed, GFP-marked stem cells are null for both *Apc* and *Apc2* (*APC^−/−^*). Over the next 21 days, many of these stem cells develop into GFP-marked, multilayered adenomas. See Figures S1B-E and S2A. **(B)** Loss of E-cad in supra-basal layers of *APC*^−/−^ tumors. E-cad∷mTomato (red hot LUT) is expressed in control tissue (top panels) and in basally localized cells within multi-layered *APC*^−/−^ clones (middle panel) but is absent from supra-basal cells (bottom panel). Each image is a single z-section of the respective conditions. **(C)** Progressive loss of E-cad∷mTomato expression in *APC*^−/−^ clones. Percentages of all *APC*^−/−^ clone cells that exhibit E-cad∷mTomato are shown for single-layered clones, basal layers of multilayered clones, and supra-basal layers of multilayered clones. 67 single-layered tumors and 38 multilayered *APC*^−/−^ clones pooled from *n* = 4 midguts at 21 days post-induction. **(D)** Experimental schema for F-H. GFP-marked, *APC*^−/−^ stem cells are generated at 1 day post-eclosion, and gene expression is manipulated specifically within these cells. At 22 days post-eclosion (21 days post-induction), the resulting, GFP-marked stem cell clones are analyzed. See Figures S2A and S2D for genetic strategy. **(E)** Domain structure of wild-type (WT) and mutant *E-cad* alleles. E-cad^dCR4h^ lacks the extracellular adhesion domain [30]. E-cad^ΔJM^ lacks the intracellular binding domain for p120-catenin [34]. **(F-H)** Ectopic expression of E-cad in nascent *APC*^−/−^ tumors inhibits tumor progression in a p120-dependent, adhesion-independent manner. Images **(F)** and cell counts **(G)** of control or *APC*^−/−^ clones with clone-autonomous expression of the indicated transgenes. Cells in clones are marked by GFP. Clone boundaries are outlined in white. In G, *n* = 3 midguts per genotype; *p-*values by Mann-Whitney *U*-test. **(H)** Frequency of multilayered clones as a percentage of total clones. *n* = 4 midguts per genotype. *p-*values by unpaired *t*-test. For **(C**, **G**, **H)**, one of three independent experiments is shown with *n* midguts per experiment as indicated. For box-and-whisker plots, boxes show median, 25th and 75th percentiles, and whiskers are minimum and maximum values. Representative images shown in each panel. All scale bars, 50 μm.

As human *APC* cancers progress, they often lose expression of E-cadherin (E-cad, also *shotgun*) [27, 28]. We therefore examined whether *Drosophila* midgut *APC*^−/−^ tumors lose *E-cad* as they develop. In control midguts, E-cad∷mTomato [29] localized prominently to lateral cell membranes, as expected (Figure 1B, S1E) [8]. In single-layered *APC*^−/−^ clones and in the basal layers of multilayered *APC*^−/−^ clones, E-cad∷mTomato was still present. However in 75% of supra-basal layers E-cad∷mTomato was not detected (Figures 1B, 1C, S1E; see Table S1 for all experimental genotypes). Thus, *Drosophila APC*^−/−^ tumors, like their human counterparts, downregulate E-cad as they progress.

Since human E-cad is an epithelial tumor suppressor, we wondered whether forced *E-cad* expression would suppress tumor formation. Hence, we assessed the tumorigenicity of *APC*^−/−^ stem cells that ectopically overexpressed *E-cad* (Figures 1D, S2D). *E-cad* overexpression had no effect on the sizes of control clones, but markedly reduced the sizes of *APC*^−/−^ clones (Figures 1F, 1G; see Tables S2, S3 for clone statistics). Furthermore, *E-cad* overexpression sharply reduced the frequency of *APC*^−/−^ multilayering, from 41.4 ± 6.3% of *APC*^−/−^ clones to only 7.1 ± 3.1% of *E-cad*-expressing *APC*^−/−^ clones (Figure 1H). Thus, *E-cad* acts as a tumor suppressor in the *Drosophila* midgut.

We sought to determine how E-cad suppresses midgut tumorigenesis. One model posits that loss of E-cad weakens cell-cell adhesion, facilitating cell invasion that is characteristic of advanced tumor stages. To test whether tumor suppression by E-cad involves adhesion, we forced *APC*^−/−^ stem cells to overexpress an adhesion-incompetent mutant that lacks extracellular adhesion motifs, E-cad^dCR4h^ (Figure 1E) [30]. E-cad^dCR4h^ substantially prevented *APC*^−/−^ clone overgrowth and multilayering (Figures 1F-H). E-cad^dCR4h^ overexpression did not alter control clone sizes (Figure 1G). These striking results demonstrate that tumor suppression by *Drosophila* E-cad does not require cadherin-mediated extracellular adhesion.

E-cad’s intracellular domain associates with two catenin-family transcription factors, β-catenin (Armadillo) and p120-catenin (p120) [31]. Although β-catenin contributes to *APC*-driven tumorigenesis in both *Drosophila* midgut and mammalian intestine [21, 22, 32, 33], it associates with both tumor suppressive alleles (*E-cad*^*WT*^ and *E-cad^dCR4h^*) and the allele E-cad^ΔJM^, which we show below is non-suppressive (Figure 1E). Thus, E-cad’s ability to suppress tumor growth cannot be attributed to β-catenin sequestration.

We next examined p120. We previously found that, during steady-state turnover [8], E-cad prevents p120 from activating transcription of the EGF protease *rhomboid* (Figure S1A), likely by sequestering p120 at the enterocyte cortex. We thus examined whether E-cad suppresses tumorigenesis by binding p120. We forced *APC*^−/−^ stem cells to express E-cad^ΔJM^, a mutant with a juxtamembrane deletion that abrogates p120 but Not β-catenin binding (Figure 1E) [34]. Unlike E-cad and Ecad^dCR4h^, E-cad^ΔJM^ failed to β suppress tumorigenesis. *APC*^−/−^ clones overexpressing *E-cad*^*ΔJM*^ grew to sizes comparable to *APC*^−/−^ clones, and a similar proportion became multilayered (Figures 1G, 1H). E-cad^ΔJM^ overexpression did not alter the sizes of control clones (Figure 1G). These results imply that E-cad-p120 binding is crucial for tumor suppression.

If E-cad sequesters p120 to suppress tumorigenesis, loss of p120 should also suppress tumorigenesis. To test this prediction, we depleted *p120* from *APC*^−/−^ stem cells using RNAi. With *p120* depletion, *APC*^−/−^ clones accumulated significantly fewer cells compared to *APC*^−/−^ clones (Figures 1F, 1G). They also exhibited less multilayering (Figure 1H). *p120* RNAi did not affect control clone sizes (Figure 1G). These findings, combined with the loss of tumor suppression by *E-cad*^*ΔJM*^, imply that downregulation of E-cad promotes tumorigenesis by dysregulating p120.

The key function of p120 during steady-state turnover is to activate *rhomboid* [8]. We therefore wondered whether p120 contributes to *APC*^−/−^ tumor development by *rhomboid* activation. First, we examined expression of a *rhomboid-lacZ* reporter (*rho-lacZ*; Figure 2A), which we built into a genetic system for generating negatively marked *APC*^−/−^ clones (Figure S2B) [24]. In this system, all cells initially express GFP, and Flp/FRT recombination generates *APC*^−/−^ stem cells that are unlabeled. Importantly, all cells possessed and were capable of expressing the *rhomboid-lacZ* transgene.

**Figure 2.**
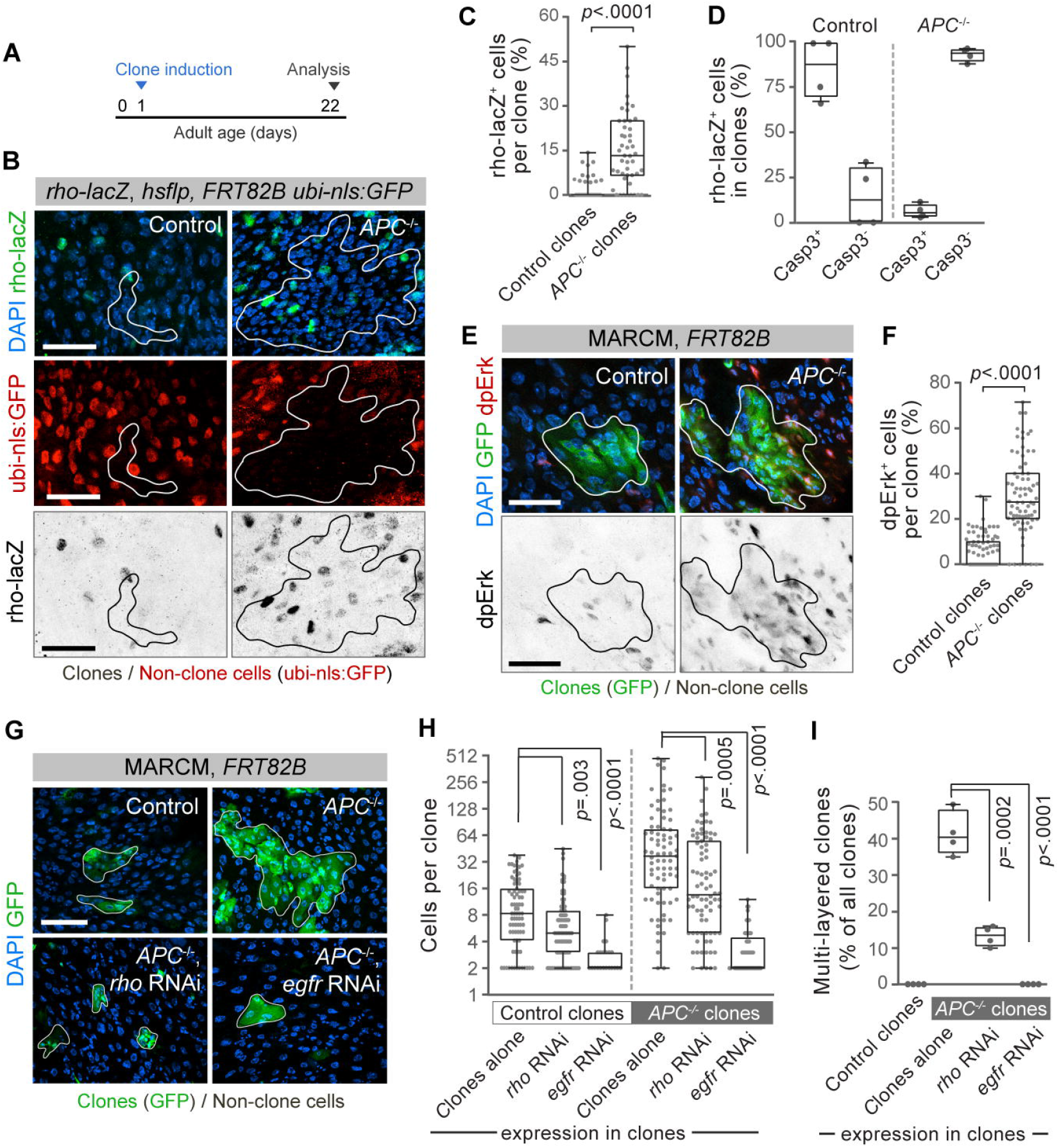
Hyperactivation of *rhomboid* and Egfr in non-apoptotic *APC*^−/−^ cells is essential for tumor growth and multilayering. **(A)** Experimental timeline for B-I. Clones are induced one day post-eclosion, and midguts are analyzed at 22 days post-eclosion (21 days post-induction). **(B-C)** Cells in *APC*^−/−^ clones express *rhomboid* more frequently than cells in control clones. The genetic schema in Figure S2B was used to generate unmarked clones in background of GFP-expressing ‘non-clone’ cells (red pseudocolor). Clone boundaries are outlined in white (top and middle rows) and black (bottom row). **(B)** Immunostaining for a *rhomboid-lacZ* reporter (*rho-lacZ*) in midguts with control clones (left column) and *APC*^−/−^ clones (right column). **(C)** Percentage of cells per clone that express *rhomboid-lacZ*. In the control dataset, 42 clones contain no *rhomboid-lacZ*^+^ cells (0%). Clones from *n* = 3 midguts per genotype; *P* values by Mann-Whitney *U*-test. **(D)** *rhomboid* expression in *APC*^−/−^ cells no longer correlates with apoptosis. The genetic schema in Figure S2B was used to generate either control or *APC*^−/−^ clones in midguts with the *rhomboid-lacZ* reporter. Midguts were immunostained for β-galactosidase (*rhomboid-lacZ*) and cleaved Caspase3. Graph shows percentages of LacZ^+^ clone cells that are Casp3^+^ or Casp3^−^. Whereas most LacZ^+^ cells in control clones are Casp3^+^, most LacZ^+^ cells in *APC*^−/−^ clones are Casp3^−^. *n* = 4 midguts per condition. See Figures S3A and S3B for representative images. **(E-F)** Cells in *APC*^−/−^ clones activate Erk more frequently than cells in control clones. **(E)** The genetic schema in Figure S2A was used to generate GFP-marked clones (green, top row). Clone boundaries are outlined in white (top row) and black (bottom row). Erk activation was assessed by immunostaining for di-phosphorylated Erk (dpErk) (top row, red; bottom row, inverted grayscale). **(F)** Percentages of cells per clone that exhibit dpErk. In the control dataset, 47 clones contain no dpErk^+^ cells (0%). Clones from *n* = 3 midguts per genotype; *p-*values by Mann-Whitney *U*-test. **(G-I)** Tumor growth and multilayering require tumor-autonomous *rhomboid* and *egfr*. **(G)** The genetic schema in Figures S2A and S2D was used to generate GFP-marked clones (green) with clone-autonomous expression of the indicated RNAi transgenes. Clone boundaries are outlined in white. **(H)** Sizes of control or *APC*^−/−^ clones that express the indicated transgenes. *n* = 3 midguts per genotype; *p-*values by Mann-Whitney *U*-test. **(I)** Frequency of multilayered clones as a percentage of total clones. *n* = 4 midguts per genotype. *p-*values by unpaired *t*-test. For **(C-D, F, H-I)**, one of three independent experiments is shown with *n* samples as specified for each experiment. For box-and-whisker plots, the boxes show median, 25^th^ and 75^th^ percentiles, and whiskers are minimum and maximum values. Representative images shown in each panel. All scale bars, 50 μm.

We observed widespread expression of *rhomboid-lacZ* in midguts containing *APC*^−/−^ tumors. After 21 days of clone development, *rhomboid-lacZ* was expressed by 15.3 ± 12.4% of cells in *APC*^−/−^ clones but only 1.7 ± 3.4% of cells in control clones (Figures 2A-C). Furthermore, in midguts that contained *APC*^−/−^ clones, *rhomboid-lacZ* was detected in 20.0 ± 12.7% of cells outside the clones (henceforth referred to as “non-clone cells”); whereas in midguts with control clones, *rhomboid-lacZ* was detected in on-ly 2.1 ± 1.7% of non-clone cells (Figures 2B, 3A). This global upregulation of *rhomboid-lacZ* in tumor-containing guts was accompanied by a pronounced increase in *rhomboid* mRNA (Figure 4F). Intriguingly, 76% of *rhomboid*-expressing non-clone cells localized within ~2 enterocyte diameters (~30 μm) of *APC*^−/−^ clones (Figures S4A, S4B). Increased *rhomboid* expression was not merely caused by *APC*^+/−^ heterozygosity of non-clone cells (*c.f.* Figure S2B) since numbers of *rhomboid-lacZ*^+^ cells in *APC*^+/−^ midguts and *APC*^+/+^ midguts were similar (Figure S4E). Thus, as *APC*^−/−^ tumors develop, *rhomboid* becomes hyper-induced both tumor autonomously and non-autonomously.

Since Rhomboid enables EGF secretion, its hyper-induction should lead to Egfr hyperactivation. Immunostaining for the activated, di-phosphorylated form of the Egfr effector Erk (dpErk) [4, 8, 22, 35], we found that 30.7 ± 17.0% of cells in *APC*^−/−^ clones exhibited dpErk, compared to 5.5 ± 7.1% of cells in control clones (Figures 2E, 2F). Non-clone cells also exhibited dpErk more frequently in midguts with *APC*^−/−^ clones compared to midguts with control clones (Figure 2E). Frequencies of *rhomboid* induction and Egfr activation were similar in single-layered and multilayered tumors (Figures S1H, S1I). Overall, these data show Egfr hyperactivation accompanies *rhomboid* hyperinduction during *APC*^−/−^ tumor formation.

Does elevated Rhomboid-Egfr signaling promote tumor development? We first investigated this possibility by examining whether developing tumors require tumor-autonomous *rhomboid*. We used MARCM to generate GFP-labeled, *APC*^−/−^ stem cells that additionally expressed *rhomboid* RNAi (Figures S2A, S2D). The clones arising from these *rhomboid* RNAi, *APC*^−/−^ stem cells were markedly smaller than those arising from *APC*^−/−^ stem cells (Figures 2G, 2H). Moreover, the vast majority of *rhomboid* RNAi, *APC*^−/−^ clones did not become multilayered (Figure 2I).

Consistent with this requirement for *rhomboid*, and similar to prior reports [4, 8, 22, 35–39], depleting *egfr* blocked both *APC*^−/−^ and control clone growth and *APC*^−/−^ clone multilayering (Figures 2G, 2H, 2I). Thus, hyper-induction of *rhomboid* in tumors promotes tumorigenesis, likely by potentiating EGFs secretion and consequent Egfr hyperactivation.

We next assessed whether tumor development requires non-autonomous *rhomboid* in non-clone cells. We specifically manipulated gene expression in non-clone cells by combining the GeneSwitch system (*GSG2326*; Figures S2C, S2E) with Flp/FRT recombination to generate *APC*^−/−^ stem cells that lack the GeneSwitch Gal4 driver [24]. In this system, oral administration of RU486 induces UAS-transgene expression specifically in non-clone cells and not *APC*^−/−^ cells. Non-clone cells are distinguished from *APC*^−/−^ cells by expression of a recombination-sensitive RFP transgene.

We found that expression of *rhomboid* RNAi in non-clone cells dramatically reduced *APC*^−/−^ clone sizes (Figure 3C), such that they approached the sizes of control clones with unmanipulated non-clone cells (Figure 3D, 3E). Multilayering was also substantially reduced (Figure 3F). Overexpressing *E-cad* or depleting *p120* had similar effects (Figures 3D-3F).

**Figure 3.**
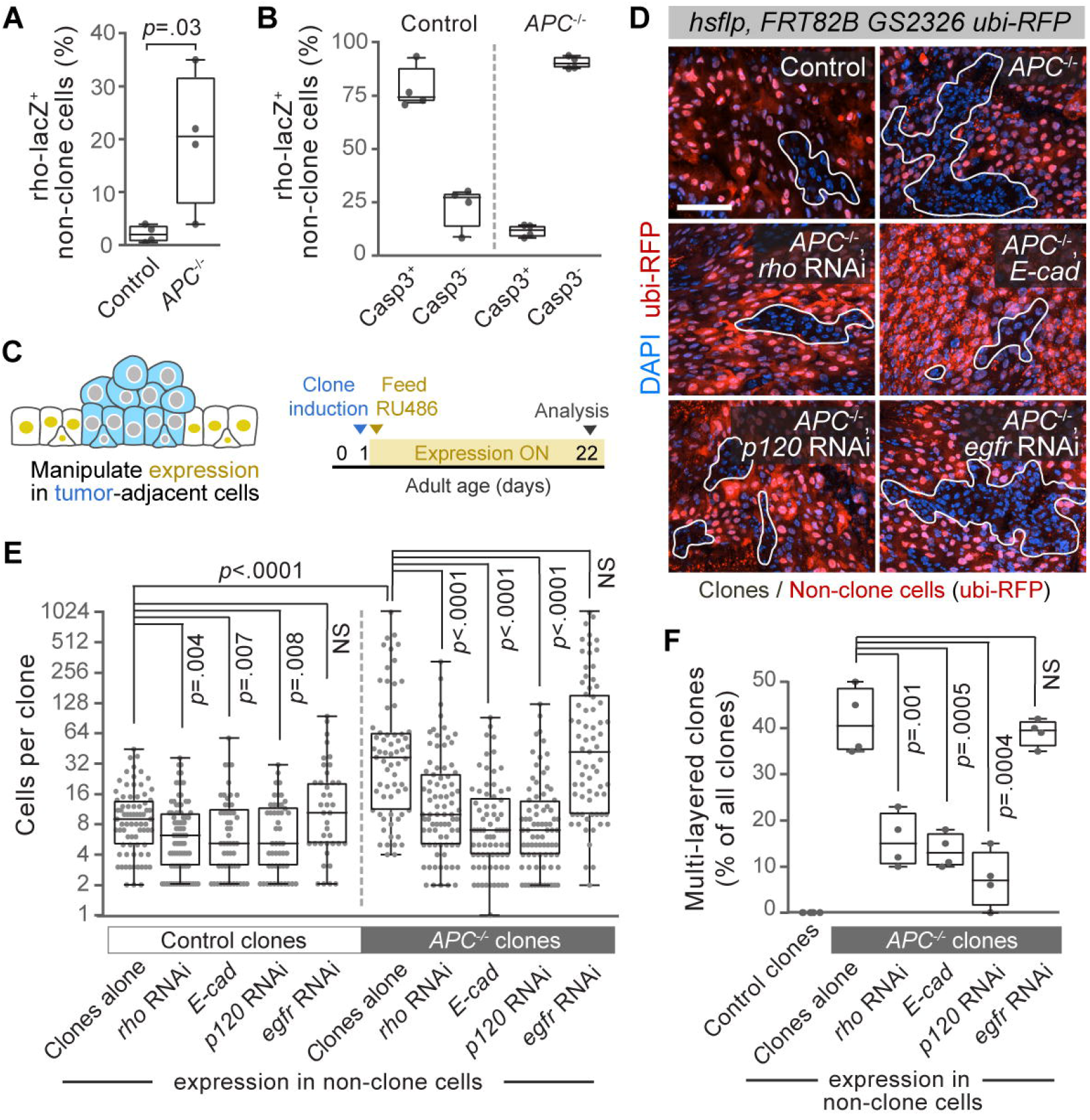
Hyperactivation of *rhomboid* occurs in non-apoptotic cells surrounding tumors and is essential for tumor growth and multilayering. **(A)** Non-clone cells that surround *APC*^−/−^ clones express *rhomboid* more frequently compared to non-clone cells that surround control clones. Control and *APC*^−/−^ midgut clones were induced using the same experimental protocol and genotypes as Figures 2A-C. Each data point shows the percentage of all non-clone cells that are *rhomboid-lacZ*^+^ in one midgut. Data points in Figures 3A and 2C were obtained from the same midguts. *n* = 4 midguts per genotype. *P* values by unpaired *t*-test. **(B)** *rhomboid* expression in non-clone cells no longer correlates with apoptosis in midguts that contain *APC*^−/−^ clones. The genetic schema in Figure S2B was used to generate either control or *APC*^−/−^ clones in midguts containing *rhomboid-lacZ*. Midguts were immunostained for *rhomboid-lacZ* and Casp3. Graph shows percentages of all *rhomboid-lacZ*^+^ non-clone cells per midgut that are apoptotic (Casp3^+^) or non-apoptotic (Casp3^−^). *n* = 4 midguts per genotype. See Figures S3A and S3B for representative images. **(C-F)** Tumor growth and multilayering require tumor non-autonomous *rhomboid* but not *egfr*. **(C)** Timeline for generation of clones and concomitant genetic manipulation of non-clone cells. The genetic schema in Figures S2C, 3B was used to generate unmarked clones surrounded by RFP-marked non-clone cells that inducibly express the indicated transgenes upon administration of RU486. All control and experimental animals received RU486 from Day 1 (clone induction) to Day 22 (analysis). **(D-E)** Images **(D)** and sizes **(E)** of control or *APC*^−/−^ clones with non-clone cell expression of the indicated transgenes. Clone boundaries are outlined in white. Scale bar, 50 μm. In **(E)**, *n* = 3 midguts per genotype; *P* values by Mann-Whitney *U*-test. **(F)** Frequency of multilayered clones as a percentage of total clones. *n* = 4 midguts per genotype. *P* values by unpaired *t*-test. For **(A-B, E-F)**, one of three independent experiments is shown with *n* numbers as specified for each experiment. For box-and-whisker plots, the boxes show median, 25^th^ and 75^th^ percentiles, and whiskers are minimum and maximum values. Representative images shown in each image panel.

The strong growth inhibition that these three non-autonomous manipulations effected on *APC*^−/−^ clones contrasted with their comparatively weak effects on control clones (Figure 3E). This difference implies that tumor non-autonomous E-cad-p120-Rhomboid dysregulation specifically fosters tumorigenesis, presumably via tumor-autonomous Egfr activation. Consistent with this notion, depleting *egfr* from non-clone cells did not affect clone sizes or multilayering (Figures 3E-3F).

Together, these results demonstrate that nascent tumors can progress only when the E-cad-p120-Rhomboid pathway is disrupted in both tumors and surrounding, non-tumor cells. This dual requirement suggests that both cell populations are needed to produce EGFs in quantities sufficient to overcome robust mechanisms of feedback control.

During normal turnover, *rhomboid* is suppressed in healthy enterocytes but induced in apoptotic enterocytes (Figure S1A) [8]. This regulatory switch forms the linchpin for tissue-level cell equilibrium by spatiotemporally coupling EGF secretion to the loss of terminally differentiated cells [8]. Given this coupling, we asked whether tumorigenic subversion of cell equilibrium involves deregulation of *rhomboid* expression. To start, we investigated whether *rhomboid*-expressing cells were apoptotic by immunostaining against cleaved Caspase3 (*c.f.* Figure S3A). In control midguts, as expected [8], a small minority of *rhomboid-lacZ*^+^ cells was non-apoptotic in both clones (14.5 ± 17.0%) (Figures 2D, S3B) and non-clone cells (22.5 ± 9.8%) (Figures 3B, S3B). In midguts with *APC*^−/−^ clones, however, the vast majority of *rhomboid-lacZ*^+^ cells was non-apoptotic in both clones (92.7 ± 3.7%) (Figures 2D, S3B) and non-clone cells (89.0 ± 2.9%) (Figures 3B, S3B). This striking finding reveals that the presence of *APC*^−/−^ tumors causes *rhomboid* to be inappropriately expressed in cells that are not undergoing apoptotic elimination.

Intriguingly, although apoptotic cells comprise a small fraction of the total cell population, Suijkerbuijk *et al*. previously found that inhibiting apoptosis of non-clone cells reduces the sizes of both *APC*^−/−^ and control clones. We thus probed the role of apoptosis by expressing the potent caspase inhibitor *p35* [40] in either *APC*^−/−^ clones (Figures S2A, S2D) or non-clone cells (Figures S2C, S2E). Consistent with prior work [24], we observed that *APC*^−/−^ clones surrounded by p35-expressing non-clone cells were smaller (Figure S3K) [8, 24]. In addition, these clones exhibited reduced levels of Egfr activation and lower frequencies of multilayering compared to *APC*^−/−^ clones with control non-clone cells (Figures S3E-S3F, S3J-S3L). By contrast, *p35* expression in *APC*^−/−^ clones had no significant effect on clone sizes, Egfr activation, or multilayering (Figures S3C-S3D, S3G-S3I). Taken together, these results suggest that tumor non-autonomous apoptosis is essential for tumor Egfr activation and growth. This finding is counterintuitive considering only 11% of *rhomboid*-expressing non-clone cells are apoptotic (Figure 3B), and the contributing mechanisms are unknown.

How do nascent tumors drive *rhomboid* hyperinduction? An attractive possibility involves Jun N-terminal kinase (JNK; also *basket*/*bsk*). JNK activation is essential for *APC*^−/−^ tumors to grow [24]. Furthermore, JNK-dependent regeneration of damaged midguts is accompanied by widespread induction of *rhomboid* [38, 41, 42]. Hence, we investigated whether *APC*^−/−^ cells co-opt their neighboring, non-clone cells into expressing *rhomboid* via JNK. We first examined the kinetics of JNK and *rhomboid* expression as tumors developed over time. We generated unmarked *APC*^−/−^ stem cells in a background of GFP-labeled non-clone cells (Figure S2B), harvested midguts at 2, 5, 10, or 21 days after clone induction (Figure 4A), and immunostained non-clone cells for activated, phosphorylated JNK (pJNK) and β-galactosidase (*rho-lacZ*; Figures 4B, S4A).

**Figure 4.**
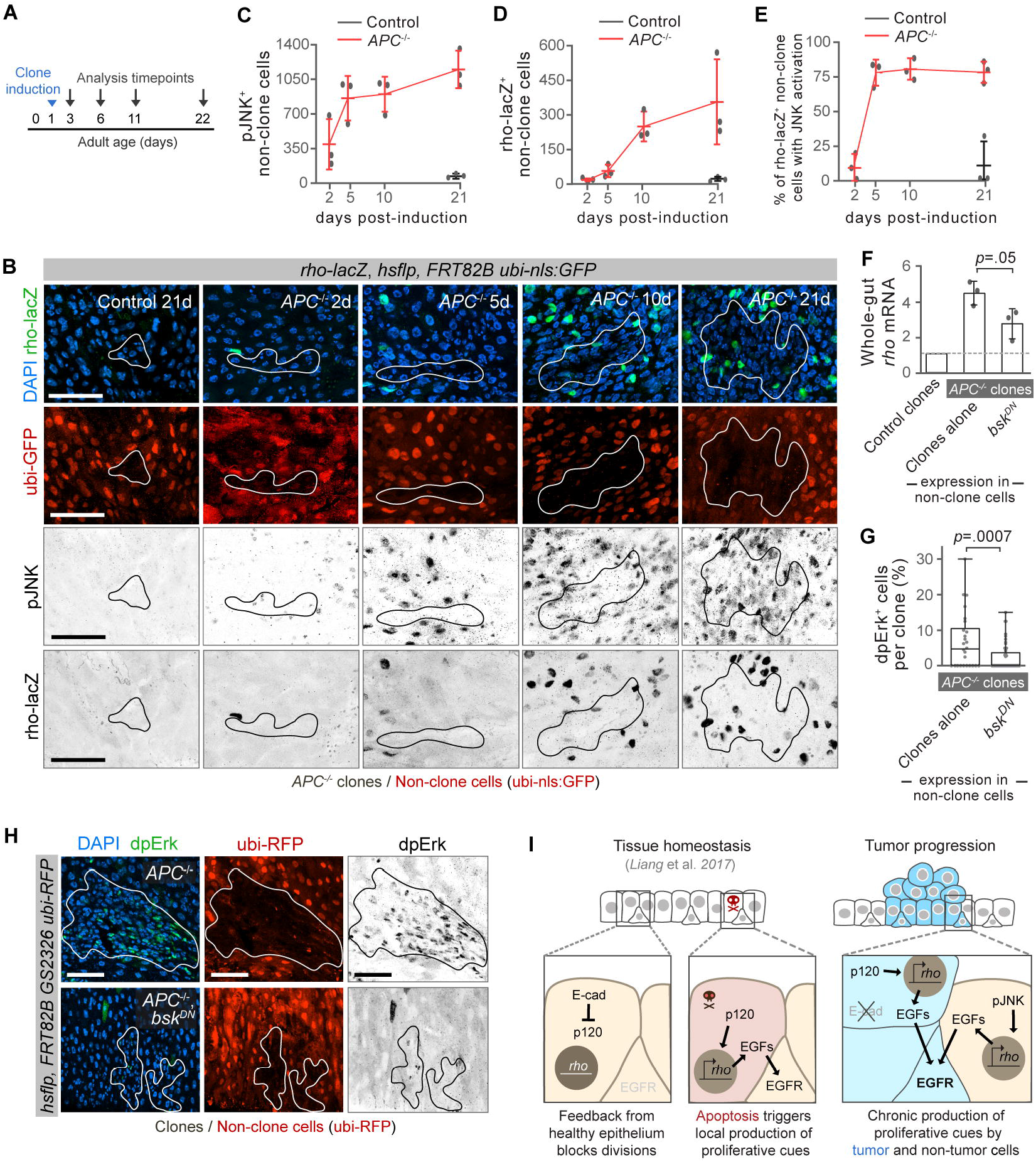
Nascent *APC*^−/−^ clones induce *rhomboid* by eliciting JNK activation in surrounding, non-tumor cells. **(A)** Experimental timeline for B-E. *APC*^−/−^ clones are induced in animals at 1 day post-eclosion and midguts are analyzed 2, 5, 10, and 21 days later. **(B-E)** Non-clone cells that surround *APC*^−/−^ clones activate JNK early in tumorigenesis and express *rhomboid* subsequently. The genetic schema in Figure S2B was used to generate unmarked control or *APC*^−/−^ clones surrounded by GFP-marked non-clone cells in midguts with *rhomboid-lacZ*. Midguts were immunostained for *rhomboid-lacZ* and phosphorylated JNK. **(B)** Representative images of midguts containing either control or *APC*^−/−^ clones at the indicated times after clone induction. Clone boundaries are outlined in white (top two rows) and black (bottom two rows). Top row shows *rhomboid-lacZ* in green. Second row shows GFP-marked non-clone cells in red. Third row shows pJNK in inverted grayscale in third row. Fourth row shows *rhomboid-lacZ* in inverted grayscale. Scale bars, 50 μm. See also Figure S4A. **(C, D)** Numbers of non-clone cells that are either pJNK^+^ **(C)** or *rhomboid-lacZ*^+^ **(D)** in midguts analyzed at the indicated times. **(E)** Most non-clone cells that express *rhomboid* also exhibit JNK activation. Graph shows the percentages of all *rhomboid*-lacZ^+^ non-clone cells that are also pJNK^+^ in midguts analyzed at the indicated times. For **(C-E)**, gray bars represent midguts with control clones and red bars represent midguts with *APC*^−/−^ clones. Each data point represents one midgut. *n* = 3 midguts per timepoint; means ± S.D. One of three independent experiments is shown. **(F-H)** JNK activation in non-clone cells promotes *rhomboid* hyper-induction and tumor cell Erk activation. The genetic strategy in Figures S2C and S2E and experimental protocol in Figure 3C was used to generate either control or *APC*^−/−^ clones and concomitantly express a dominant negative allele of JNK (*bsk^DN^*) in non-clone cells. **(F)** Inhibition of JNK in non-clone cells reduces levels of *rhomboid* mRNA. Whole-midgut qPCR was performed using midguts of the indicated genotypes. mRNA levels are shown normalized to midguts that contain control clones (left bar). Bars represent means ± S.D.; three biological replicates per condition. *P* values by unpaired *t*-test. **(G-H)** Inhibition of JNK in non-clone cells reduces Erk activation in *APC*^−/−^ clones. Immunostaining for dpErk was performed on *APC*^−/−^ clone-containing midguts with or without *bsk*^*DN*^ expression in non-clone cells. Percentages of cells per clone that are dpErk^+^ **(G)** and representative images **(H)** are shown. In the *bsk*^*DN*^ condition, 35 clones contain zero dpErk^+^ cells (0%). Clones from *n* = 3 midguts per genotype; *P* values by Mann-Whitney *U*-test. One of three independent experiments is shown. Scale bars, 50 μm. **(I)** Model. Tumor establishment requires that tumorigenic stem cells de-stabilize cell equilibrium by coercing non-apoptotic cells to express *rhomboid*. During healthy turnover (left), stem cell division is coupled to enterocyte death because expression of *rhomboid*, and hence secretion of mitogenic EGFs, occurs via apoptotic downregulation of E-cad and consequent release of p120-catenin [8]. For new tumors to become established (right), tumor-initiating stem cells decouple division from death by instigating widespread expression of *rhomboid* in cells that are not apoptotic. *rhomboid* hyper-induction is initially tumor non-autonomous, via activation of JNK in non-clone cells. It subsequently becomes tumor autonomous, via downregulation of E-cad and consequent activity of p120-catenin in tumor cells.

We found that numbers of JNK-activated non-clone cells surged rapidly and dramatically during *APC*^−/−^ tumorigenesis (Figures 4B, 4C). pJNK^+^ non-clone cells climbed sharply from 2-5 days and remained extremely high from 5-21 days. At 21 days, pJNK^+^ non-clone cells were markedly elevated (1151.0 ± 190.2 pJNK^+^ cells) compared to either wild-type guts with control clones (10.5 ± 8.4) or genotype-matched *APC*^+/−^ guts in which *APC*^−/−^ clones had not been induced (35.0 ± 9.1) (Figures 4B, S4D). Thus, *APC*^−/−^ clones prompt non-clone cells to acutely hyperactivate JNK early in tumorigenesis, before the mutant clones have become multilayered tumors (Figure S1F).

JNK activation preceded *rhomboid* induction (Figures 4B-D). After 2 days of *APC*^−/−^ clone development, *rhomboid-lacZ*^+^ non-clone cells were still at control levels. After 5 days, they had increased only slightly. From 5-10 days, however, *rhomboid-lacZ*^+^ cells increased dramatically, and from 10-21 days they remained highly elevated. In these tumor-containing guts, 78.3 ± 7.6% of non-clone cells that had turned on *rhomboid* were also pJNK^+^ (Figures 4B, 4E, S4A). By contrast, in *APC*^+/−^ guts lacking *APC*^−/−^ clones, only 9.8 ± 3.2% of *rhomboid*-expressing cells were also pJNK^+^ (Figure S4F). Co-localization of *rhomboid* expression and activated JNK in the same non-clone cells (Figure 4E), together with the general delay in *rhomboid* activation relative to JNK (Figures 4B-4D), raise the possibility that JNK induces *rhomboid*.

To investigate this possibility, we concomitantly generated *APC*^−/−^ clones and inhibited JNK in non-clone cells via RU486-inducible expression of *bsk*^*DN*^ [43, 44] (Figures S2C, S2E). This manipulation reduced *rhomboid* mRNA by 40% and diminished Egfr-activated tumor cells by 74% (Figures 4F-4H). Furthermore, *APC*^−/−^ clones were substantially smaller, as also observed by Suijkerbuijk *et al.* [24], and exhibited dramatically less multilayering (Figures S3J-3L).

By comparison, tumor-autonomous expression of *bsk*^*DN*^ in *APC*^−/−^ clones did not significantly reduce clone sizes or affect multilayering (Figures S3G-3I), possibly reflecting that tumor cells do not activate JNK until later stages (Figure 4B). In control midguts, expressing *bsk*^*DN*^ in clones or in surrounding non-clone cells did not change clone sizes (Figures S3H, S3K), consistent with prior work [45]. Combined, these kinetic and functional analyses imply a model in which *APC*^−/−^ cells acutely hyperactivate JNK in non-clone cells, which consequently induce *rhomboid* to activate Egfr and promote tumor growth.

## Discussion

During steady-state turnover of the *Drosophila* midgut, expression of the EGF protease *rhomboid* is suppressed in healthy enterocytes and activated in enterocytes undergoing apoptotic elimination [8]. This mechanism provides feedback control so that the mitogenic EGFs Spitz and Keren, both regulated by Rhomboid, become available specifically at the time and place that replacement cells are needed [8]. Here, we have shown that nascent tumors transform this feedback control into feed-forward activation by instigating widespread induction of *rhomboid*, including in non-apoptotic cells (Figure 4I). This inappropriate *rhomboid* induction enables EGFs to be secreted chronically, which in turn drives production of new cells regardless of tissue need.

Feedback EGF signaling is transformed into feed-forward activation via sequential non-autonomous and autonomous mechanisms that effectively short-circuit the homeostatic pathway for *rhomboid* activation. Even before *APC*^−/−^ cells manifest as tumors, they induce *rhomboid* in wild-type neighbor cells via non-autonomous activation of JNK. During subsequent growth, tumors autonomously activate *rhomboid* via loss of E-cad and release of p120-catenin in an apoptosis-independent manner. R*homboid* dysregulation in both tumor and non-tumor cells is required to form multilayered adenomas. This dual requirement suggests that high levels of EGFs are necessary to overcome robust enforcement of cell equilibrium.

Suijkerbuijk and colleagues elegantly demonstrated that competition between *APC*^−/−^ cells and wild-type cells leads to tumor growth [24], but the growth-promoting mechanism remained unknown. We suggest cell competition may promote growth by deregulating *rhomboid*, which would lead to consequent activation of Egfr. If so, a major implication is that tumor/non-tumor cell competition acts by directly subverting pathways that mediate normal homeostasis. Whether cell competition during mammalian tumorigenesis [46–51] follows a similar template will be important to determine.

The mechanisms that enable establishment of *Drosophila APC*^−/−^ tumors may illuminate initiation of human colorectal cancers, which are tightly associated with *APC* inactivating mutations. Intriguingly, Rhomboids, E-cad, and EGFR have been implicated in human tumor progression [13, 17, 52–54], suggesting this signaling axis may be conserved. This possibility confers particular interest on two of our findings. First, while loss of E-cad is canonically thought to promote metastasis via loss of cell-cell adhesion, we uncovered a crucial role during early-tumor development: dysregulation of p120-catenin to drive EGF signaling. Whether a similar relationship between p120-catenin and EGF exists in colorectal cancer merits examination. Second, while studies of mammalian Rhomboids have focused on advanced cancers, we find *rhomboid* induction is a tumor-initiating event. Hence, examining Rhomboids in early-stage mammalian tumorigenesis may be fruitful. Overall, understanding how nascent tumors destabilize cell equilibrium may suggest strategies for preventing potentially tumorigenic cells from establishing tumors.

## Acknowledgements

S.N. was supported by a Stanford Bio-X Undergraduate Fellowship. J.L. was supported by NSF GRFP DGE-114747 and NIH T32GM007276. This work was supported by ACS RSG-17-167-01-DDC, NIH R01GM116000-01A1, and a Stanford VPUE Faculty Grant to L.E.O. Confocal microscopy was performed at the Stanford Beckman Cell Sciences Imaging Facility (NIH 1S10OD01058001A1). We thank D. Bilder for the gift of cCas-3 antibody; the Developmental Studies Hybridoma Bank for other antibodies; M. Peifer, E. Piddini, M. Fuller, H. Jiang, the Bloomington *Drosophila* Stock Center (NIH P40OD018537), the TRiP at Harvard Medical School (NIH/NIGMS R01-GM084947), and the Vienna *Drosophila* Resource Center (http://stockcenter.vdrc.at/control/main) for fly stocks; M. Mirvis, L.J. Koyama, and E.N. Sanders for helpful discussions; J.M. Knapp for writing assistance; and J. Cordero and K. Campbell for comments on the manuscript.

## Author Contributions

J.L., S.N., and L.E.O. designed the study. J.L. and L.E.O. wrote the manuscript. S.N. and Y.H.S. performed genetic crosses, tissue dissection and immunostaining. J.L. designed the experiments, built fly lines, performed qPCR experiments, quantitative analysis of microscopy data, and statistical analysis. S.N., Y.H.S., and J.L. performed confocal microscopy on tissues.

## STAR Methods

### Lead Contact and Materials Availability

Further information and requests for resources and reagents should be directed to and will be fulfilled by the Lead Contact, Lucy Erin O’Brien. This study did not generate new unique reagents.

### Experimental Model and Subject Details

Adult female flies (*Drosophila melanogaster*) were used in all experiments. Crosses and adult flies were raised at 25°C in vials containing molasses-cornmeal. Unless specified otherwise, flies were heat-shocked 1 day after eclosion to induce clones and collected 21 days after induction for dissection/ immunostaining. See Table S1 for full list of experimental genotypes.

#### Fly stocks

The following stocks were obtained from the Bloomington Stock Center: *y w shg*^*mTomato*^, *UAS-shg*, *UAS-shg*^*ΔJM*^, *UAS-egfr* RNAi (TRiP.HMS05003), *UAS-rho* RNAi (TRiP.HMS02264), *UAS-bsk*^*DN*^, and *UAS*-*p35*. *UAS-p120* RNAi (KK113572) was obtained from the Vienna *Drosophila* Resource Center. The following stocks were generous gifts: *FRT82 APC2*^*G10*^ *APC1*^*Q8*^ (from M. Peifer), *hsflp*^*122*^; *FRT82 ubi-GFP* and *hsflp*^*122*^; *FRT82 GS2326 ubi-RFP* (from E. Piddini [24]), *UAS-shg*^*dCR4h*^ (from M. Fuller), and *rho*^*X81*^(*rhomboid-lacZ*, from H. Jiang). Other stocks (from our previous studies [8,55]): *w; FRT82* and *w UAS-CD8:GFP hsflp*^*122*^; *tubGAL4; FRT82 tubGAL80*. Detailed information on *Drosophila* genes and stocks is available from FlyBase (http://flybase.org/).

### Method Details

#### Induction of stem cell clones

Tumor clones were generated using three separate labeling systems (Figure S2). For all three labeling systems, tumor clones were generated by collecting adult flies one day post-eclosion and performing two 30-min, 38.5°C heat shocks separated by a 8-min chill on ice. Flies were returned to 25°C until time of dissection. For experiments which manipulated gene expression in adjacent tissue after tumor induction (Figure S2C; also known as the “pLoser” system [24]), both control and experimental flies were fed RU486 upon returning to 25°C post-heat shock until time of dissection (see “GeneSwitch induction” below).

#### GeneSwitch induction

To induce expression of the GeneSwitch driver, *GS2326*, adult flies (control and experimental cohorts) were fed RU486. RU486 (Sigma-Aldrich) was dissolved in dH_2_O to reach a working concentration of 25 μg/mL. This solution was used to prepare yeast paste, which was fed to flies as a supplement to their standard cornmeal–molasses diet for the duration of induced gene expression. Drug-containing yeast paste was replenished every three days.

#### Bleomycin feeding

Bleomycin (Sigma-Aldrich) was prepared at a working concentration of 25 μg/ml dissolved in ddH_2_O with 5% sucrose. This solution was used to prepare yeast paste, which was fed to flies as a supplement to their standard cornmeal–molasses diet for 5 hours.

#### Immunohistochemistry and microscopy

Immunohistochemistry samples were prepared by incubating in fixative (8% formaldehyde, 200 mM Na cacodylate, 100 mM sucrose, 40mM KOAc, 10mM NaOAc, and 10mM EGTA) for 20 min at room temperature. Fixed issues were immunostained and mounted in agarose (see also [8, 55]). Anti-GFP and anti-RFP antibodies were used to improve detection of *ubi-GFP* and *ubi-RFP* expression in tumor labeling systems (Figure S2). Anti-RFP was used to detect *shg*^*mTomato*^. Primary antibodies: mouse anti-β-galactosidase (1:400, Promega Z3781), rabbit anti-cleaved caspase-3 (1:400, Cell Signaling, gift from D. Bilder), rabbit anti-dpErk (1:200, Cell Signaling 4370P), mouse anti-Coracle (1:400, DSHB C615.16), mouse anti-Discs large (1:400, DSHB 4F3), rabbit an-ti-pJNK pTPpY (1:500, Promega V7931), chicken anti-GFP (1:400, Invitrogen A10262) rabbit anti-RFP (1:500, Invitrogen R10367), and mouse anti-RFP (1:500, Invitrogen RF5R). Secondary antibodies: Alexa Fluor 488-, 555-or 647-conjugated anti-rabbit, anti-mouse, or anti-chicken IgGs (1:800, LifeTechnologies A31570, A11001, A11039, A32728, A32732, and A21244). Nuclei were stained with DAPI (LifeTechnologies, 1:1,000). Samples were mounted in ProLong (LifeTechnologies). Imaging of samples was performed on a Leica SP8 confocal microscope, with serial optical sections taken at 3.5 μm intervals through the entirety of whole-mounted, immunostained midguts.

#### qRT-PCR

mRNA was extracted from whole midguts using Trizol reagent (Invitrogen); four midguts per biological replicate. Extracted RNA was used for cDNA synthesis with Invitrogen SuperStrand III First Script Super Mix (Invitrogen). Real-time PCR was performed using the relative standard curve method with SYBR GreenER Supermix (Invitrogen) on a StepOnePlus ABI machine. Each biological replicate was assessed in three technical replicate experiments. Expression levels were normalized to non-tumorous midguts expressing control clones; *rp49* transcripts were used as a reference. Primers were from [8, 56]. Primer sequences from 5’ to 3’ – *rp49* Fwd: CGGATCGATATGCTAAGCTGT, *rp49* Rev: CGACGCACTCTGTTGTCG, *rhomboid* Fwd: GAGCACATCTACATGCAACGC, and *rhomboid* Rev: GGAGATCACTAGGATGAACCAGG.

#### Study design

Sample sizes were chosen based on our previous studies [8, 55], which also characterized changes in clone sizes and midgut cell numbers; see also Table S2. Most experiments were replicated three times; see respective figure legends. No exclusion criteria were applied. No sample randomization or blinding was performed.

### Quantification and Statistical Analysis

#### Clone visualization and quantification

Tumors were visualized (1) as z-stacks using Fiji [57] and (2) in 3D using the Bitplane Imaris software. For each midgut, all clones within the R4 and R5 regions [58] were analyzed (see Figure S1B). Cells per tumor were measured as the number of DAPI^+^ nuclei within the labeled clone boundary (as determined by the presence or absence of respective labeling proteins). All clone counts were performed manually. To categorize clones as single-layered or multilayered, each clone was viewed from the sagittal plane in Bitplane Imaris. Clones were designated as single-layered if all polyploid enterocytes possessed both (1) a free luminal surface that was not juxtaposed to other cells or to presumptive basement membrane/visceral muscle and (2) a basal surface that was not juxtaposed to other cells but was juxtaposed to presumptive basement membrane/visceral muscle. Clones were designated as multi-layered if one or more polyploid enterocytes lacked either or both of these criteria. Statistical parameters for clone reported in Table S3.

#### Statistical analysis

All statistical analyses were performed using Graphpad Prism 7. For comparisons of clone size distributions, unpaired two-tailed Mann–Whitney *U*-tests were used to assess statistical significance. To compare the frequencies of mutilayered clones, cell numbers or percentages, and mRNA levels, unpaired two-tailed *t*-tests were used to assess statistical significance. No methods were applied to test the assumptions for respective statistical approaches. Statistical parameters - e.g. sample sizes (*n*) and values represented by error bars or boxplots - are reported in respective figure legends; *p-*values are reported in respective graphs. For each experiment, n represents the number of midguts per condition.

### Data and Code Availability

This study did not generate or analyze any datasets/code.

