## Supplemental Materials for "Tumor establishment requires tumor autonomous and non-autonomous deregulation of homeostatic feedback control"

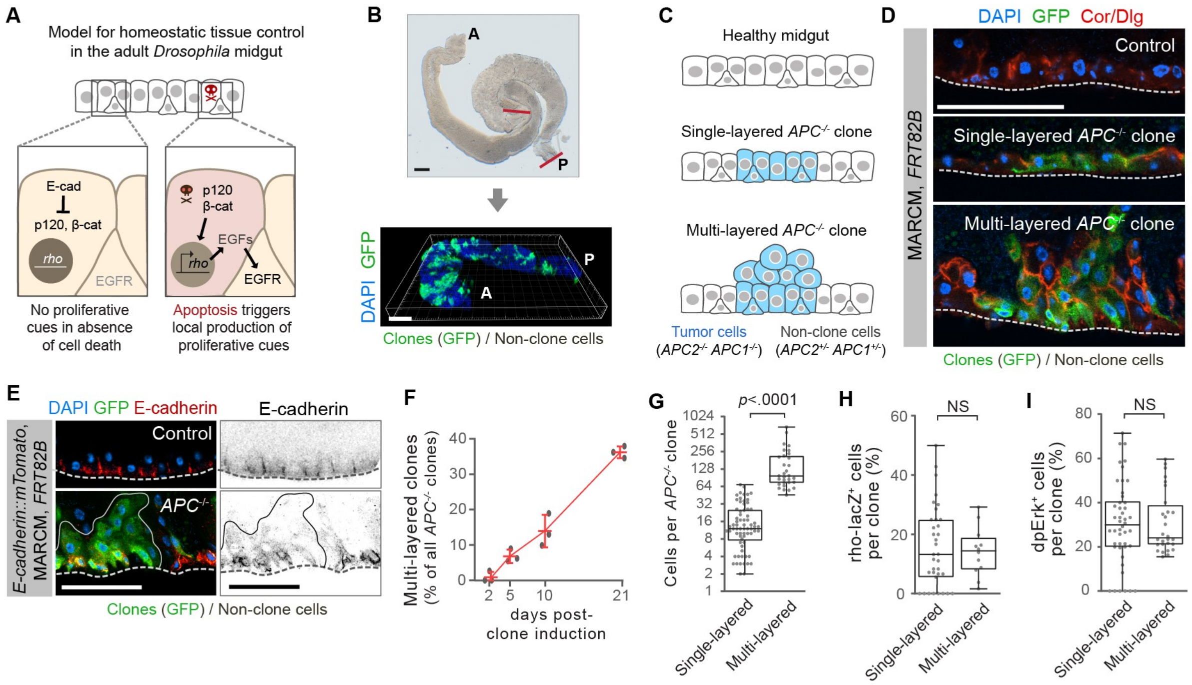


**Figure S1. Analysis of *APC*$^{-/-}$ clones in the adult *Drosophila* midgut. Related to Figure 1.**

**(A)** Cell equilibrium in the *Drosophila* adult midgut arises through coupling of stem cell divisions to enterocyte apoptosis. (Left) In the absence of enterocyte apoptosis, stem cells lack essential EGF mitogens and are therefore quiescent. Healthy enterocytes do not secrete EGFs because enterocyte E-cad prevents p120-catenin and β-catenin from activating transcription of the obligate EGF protease *rhomboid*. (Right) During physiological apoptosis, enterocytes downregulate E-cad. Loss of E-cad enables p120- and β-catenin to activate transcription of *rhomboid*. Rhomboid cleaves EGFs for secretion; the secreted EGFs activate nearby stem cells for replacement divisions. Cartoon adapted from [S1].

**(B)** Midgut regions R4 and R5 were used for clonal analyses. (Left) Brightfield image of whole mount midgut. The R4-R5 regions [S2] are demarcated by red lines. (Right) 3D reconstruction of R4 and R5 regions from a midgut that contains 21-day, GFP-labeled, *APC*$^{-/-}$ clones (green). Clone size quantifications were performed by digitally isolating R4 and R5 in Fiji and visualizing the 3D reconstructed organ in Imaris. A=anterior, P=posterior. DAPI (blue) labels nuclei.

**(C-D)** Cellular organization of *APC*$^{-/-}$ clones. Cartoons **(C)** and images **(D)** show sagittal views of control tissue (top), a representative single-layered *APC*$^{-/-}$ clone (middle), and a representative multilayered *APC*$^{-/-}$ clone (bottom). Single-layered *APC*$^{-/-}$ clones resemble control tissue. Multilayered clones creates adenomatous masses that protrude into the midgut lumen. In **(D)**, immunostaining (red) for Coracle (Cor) and Discs large (Dlg) reveals cell outlines, GFP (green) labels *APC*$^{-/-}$ clones, and DAPI (blue) labels all nuclei. Clones are 21 days post-induction.

**(E)** Saggital views of E-cad. E-cad::mTomato (red hot LUT) is expressed in control tissue (top panels) and in basally localized cells within multilayered APC$^{-/-}$ clones (bottom panels; GFP, green) but is absent from supra-basal cells. Dotted line indicates the basement membrane.

**(F)** *APC*$^{-/-}$ clones progress from single-layered to multilayered over time. Graph shows the percent of all *APC*$^{-/-}$ clones that exhibit multilayering at 2, 5, 10, and 21 days after clone induction. Each data point represents one midgut. *n* = 3 midguts per timepoint; means ± S.D are shown.

**(G-I)** Comparison of single-layered and multi-layered *APC*$^{-/-}$ clones by clone size **(G)**, percentages of cells per clone that express *rhomboid-lacZ* **(H)**, and percentages of cells per clone that exhibit dpErk **(I)**. Clones from n = 3 midguts per genotype; P values by Mann-Whitney U-test. Representative images shown in each image panel. For box-and-whisker plots, the boxes show median, 25$^{th}$ and 75th percentiles, and whiskers are minimum and maximum values. Scale bars, 50 μm.

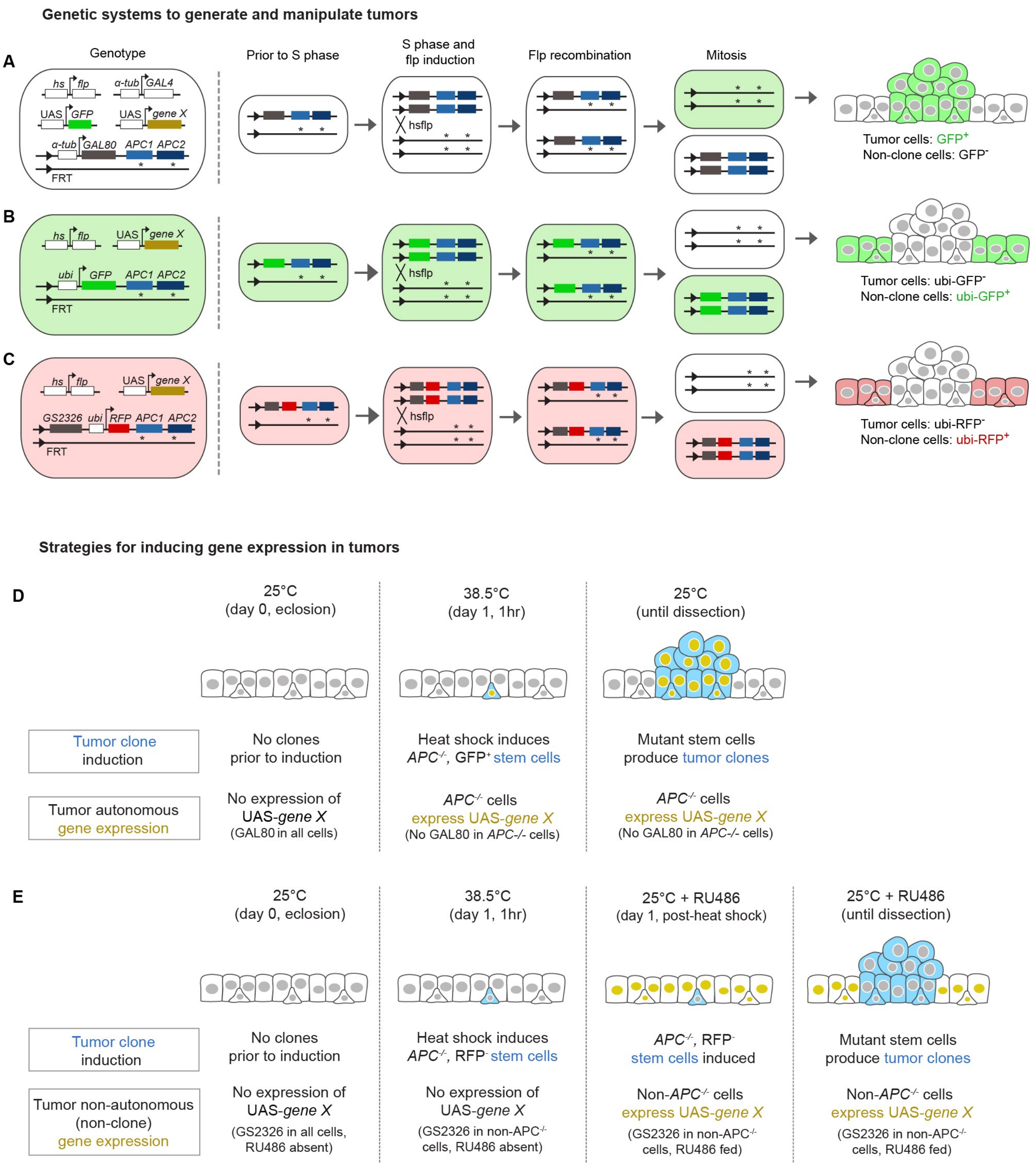


**Figure S2. Three genetic systems to generate *APC*$^{-/-}$ clones and manipulate gene expression in either clones or non-clone cells. Related to Figures 1, 2, 3 and 4.**

**(A-C)** In all three systems, all midgut cells are initially unlabeled and heterozygous for null alleles of *Apc1* (*Apc1*$^{Q8}$) and *Apc2* (*Apc2*$^{G10}$). Heat shock induction (two 30-min, 38.5°C heat shocks separated by an 8-min chill on ice) is used to activate expression of *flp*$^{122}$ recombinase, which catalyzes recombination in mitotic stem cells to generate a daughter cell that is homozygous for both *Apc1*$^{Q8}$ and *Apc2*$^{G10}$ (*APC*$^{-/-}$).

**(A)** Clone-autonomous labeling and manipulation. Schematic shows MARCM system to generate GFP-labeled *APC*$^{-/-}$ clones with concomitant genetic manipulation within clones. *Flp* recombination both generates an *APC*$^{-/-}$ daughter cell and eliminates *GAL80* specifically from that daughter. Loss of *GAL80* enables GAL4-driven expression of *UAS-GFP* and other *UAS*-transgenes (*UAS-gene X*) in the *APC*$^{-/-}$ cell and its progeny.

**(B)** Clone non-autonomous ("non-clone cell") labeling. Schematic shows generation of unlabeled *APC*$^{-/-}$ clones within a background of GFP-labeled, non-clone cells. All midgut cells initially express *ubi-GFP*. Heat-shock induction of *flp* recombinase produces an *APC*$^{-/-}$ daughter cell that lacks *ubi-GFP*; the *APC*$^{-/-}$ cell and any progeny it produces are thus unlabeled. Non-clone cells continue to express *ubi-GFP*.

**(C)** Clone non-autonomous ("non-clone cell") labeling and manipulation. Schematic shows the GeneSwitch 2326 system (also known as the "pLoser system" [S3]), which generates unlabeled *APC*$^{-/-}$ clones with concomitant labeling and manipulation of non-clone cells. Upon oral administration of RU486, *GeneSwitch2326* (*GS2326*) [S4] drives expression of *UAS*-transgenes (*UAS-gene X*) in midgut stem cells, enteroblasts, and enterocytes. Midgut cells initially express *ubi*-RFP and possess *GS2326*. Heat-shock induction of *flp* recombinase produces an *APC*$^{-/-}$ daughter cell that lacks *ubi-RFP* and *GS2326*. Non-clone cells, which continue to express *ubi-RFP*, are capable of RU486-inducible expression of *UAS*-transgenes, whereas unlabeled *APC*$^{-/-}$ cells are not. In all GeneSwitch experiments, RU486 was administered to both control and experimental cohorts.

**(D-E)** Experimental protocols for induction of *APC*$^{-/-}$ clones and concomitant genetic manipulation.

**(D)** Clone-autonomous genetic manipulation using MARCM. Animals eclose at 25ºC. Initially, all *UAS*-transgenes are silent due to ubiquitous expression of *tubGAL80*. At 1 day after eclosion, heat shock-induced *flp*$^{122}$ generates *APC*$^{-/-}$ mutant stem cells that have permanently lost *tubGAL80*. Loss of *tubGAL80* enables *tubGAL4*-driven expression of *UAS-GFP* and other *UAS*-transgenes (*UAS-geneX*) within *APC*$^{-/-}$ stem cells and any *APC*$^{-/-}$ stem cell progeny that are produced during the subsequent 25ºC chase. See also panel A.

**(E)** Clone non-autonomous (“non-clone cell” genetic manipulation using “pLoser” [S3]. Animals eclose at 25ºC. Initially, *ubiRFP* is expressed in all cells, and all *UAS*-transgenes are silent because *GeneSwitch2326* (*GS2326*), which drives expression in enterocytes, enteroblasts, and stem cells, is inactive in the absence of RU486. At 1 day after eclosion, heat shock-induced *flp*$^{122}$ generates *APC*$^{-/-}$ mutant stem cells that have permanently lost both *ubi-RFP* and *GS2326* (*c.f.* panel C), while non-clone cells retain both *ubi-RFP* and *GS2326*. To activate expression of *UAS*-transgenes (*UAS-geneX*) in these RFP-expressing non-clone cells, RU486 is orally administered immediately after heat shock and throughout the subsequent 25ºC chase. By contrast, the unlabeled *APC*$^{-/-}$ stem cells and their progeny are unresponsive to RU486. In all experiments using the “pLoser” systemn, RU486 was administered to both control and experimental cohorts.

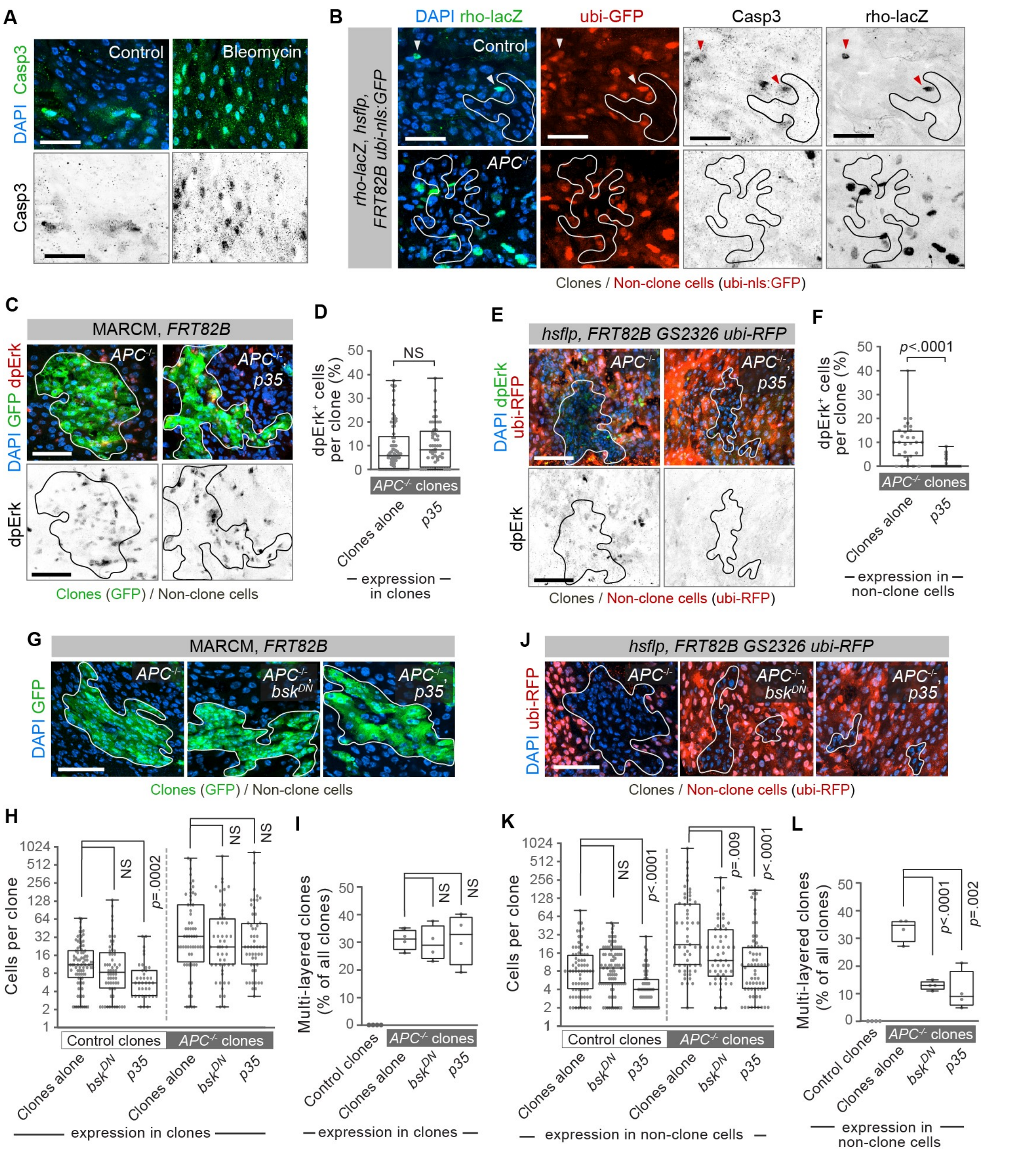


**Figure S3. Effects of inhibiting apoptosis or JNK signaling on *rhomboid* expression, Erk activation, and tumor progression. Related to Figures 2 and 3.**

**(A)** Positive control for immunostaining of apoptotic cells by anti-cleaved Caspase3 antibody. To induce apoptosis of midgut cells, the damaging agent bleomycin was administered orally. Substantially more Casp3$^{+}$ cells are visible in bleomycin-treated midguts compared to untreated control midguts.

**(B)** Sample images of midguts analyzed for Figures 2D and 3B. The genetic schema in Figure S2B was used to generate either control clones (top) or *APC*$^{-/-}$ clones (bottom) in midguts containing *rhomboid-lacZ*. Clones are unmarked, and non-clone cells are marked with RFP (red in second panel from left). Midguts were immunostained for *rhomboid-lacZ* (anti-β-galactosidase antibody; green in left panel, inverted grayscale in right panel) and Casp3 (inverted grayscale in second panel from right). In guts with control clones, most *rhomboid-lacZ*$^{+}$ cells are Casp3$^{+}$ (red arrowheads). In guts with *APC*$^{-/-}$ clones, most *rhomboid-lacZ*$^{+}$ cells are Casp3$^{-}$.

**(C-F)** Tumor autonomous activation of Erk decreases following p35-mediated inhibition of apoptosis in non-clone cells but not tumor cells. Representative images **(C, E)** and quantitation **(D, F)** of dpErk immunostaining in *APC*$^{-/-}$ clones. In **(C, D)**, *p35* was expressed in GFP-marked *APC*$^{-/-}$ clones using the genetic schema in Figures S2A and S2D. In **(E, F)**, *p35* was expressed in RFP-marked non-clone cells surrounding unmarked *APC*$^{-/-}$ clones using the genetic schema in Figures S2C and S2E. In **(F)**, 54 clones in the *p35* condition contain zero dpErk+ cells (0%).

**(G-L)** Tumor progression is reduced by inhibition of either apoptosis or JNK in non-clone cells but not tumor cells. Representative images **(G, J)** and quantitation of clone sizes **(H, K)** and multilayering **(I, L)**. In **(G-I),** the genetic schema in Figures S2A and S2D was used to express the indicated transgenes in GFP-marked control or *APC*$^{-/-}$ clones. In **(J-L)**, the genetic schema in Figures S2C and S2E was used to express the indicated transgenes in RFP-marked non-clone cells surrounding either control or *APC*$^{-/-}$ clones.

For **(C-L)**, genetic manipulations within clones were performed as in Figures 1D and genetic manipulations in non-clone cells were performed as in Figure 3C; one of three independent experiments is shown. In **(D, F, H, K)**, $n$ = 3 midguts per genotype; $P$ values by Mann-Whitney $U$-test. In **(I, L)**, $n$ = 4 midguts per genotype; $P$ values by unpaired $t$-test. Representative images shown in all panels; clone boundaries are outlined. For box-and-whisker plots, the boxes show median, 25$^{th}$ and 75th percentiles, and whiskers are minimum and maximum values. Scale bars, 50 µm.

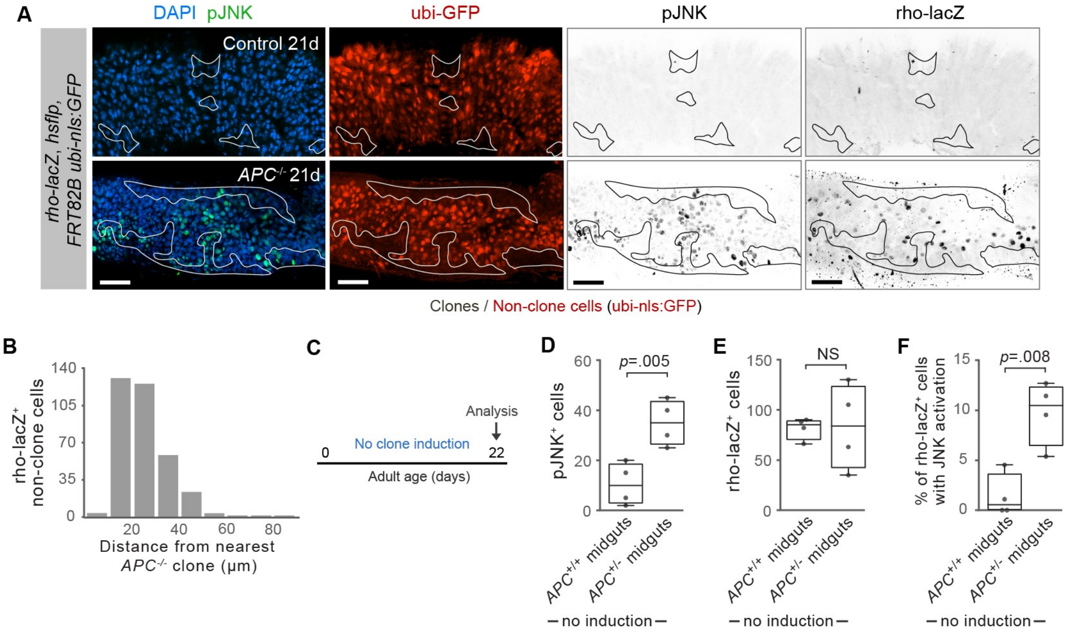


**Figure S4. JNK and *rhomboid* hyper-activation are specifically associated with *APC*$^{-/-}$ tumors. Related to Figure 4.**

**(A)** Wide-field, representative images of midguts in Figure 4B show either control or *APC*$^{-/-}$ clones at 21 days post-induction. Clone boundaries are outlined in white (left panels) and in black (right panels). pJNK$^{+}$ cells and *rhomboid-lacZ*$^{+}$ non-clone cells localize to the vicinity of *APC*$^{-/-}$ clones. Scale bars, 50 µm.

**(B)** Distances between *rhomboid-lacZ*$^{+}$ non-clone cells and the nearest *APC*$^{-/-}$ clone. Distances were measured from the center of each *rhomboid-lacZ*$^{+}$ non-clone cell nucleus to the center of the closest nucleus within the nearest clone was measured. $n$ = 344 *rhomboid-lacZ*$^{+}$ cells pooled from 3 midguts.

**(C-F)** Baseline frequencies of JNK and *rhomboid* activation in *APC*$^{+/+}$ and *APC*$^{+/-}$ midguts without clone induction. **(C)** Midguts of genotypes used in the Figure S2B schema were harvested at 22 days post-eclosion without clone induction and immunostained for *rhomboid-lacZ* and pJNK. **(D)** pJNK$^{+}$ cells are elevated in *APC*$^{+/--}$ midguts compared to *APC*$^{+/+}$ midguts but are 250-fold less abundant compared to *APC*$^{+/-}$ midguts that contain *APC*$^{-/-}$ clones (*c.f.* Figure 4C). **(E)** *rhomboid-lacZ*$^{+}$ cells are comparable in *APC*$^{+/+}$ and *APC*$^{+/-}$ midguts and substantially less abundant compared to *APC*$^{+/-}$ midguts that contain *APC*$^{-/-}$clones (*c.f.* Figure 4D). **(F)** Percentage of all *rhomboid*-lacZ$^+$ midgut cells that are also pJNK$^+$ is elevated in *APC*$^{+/-}$ midguts compared to *APC*$^{+/+}$ midguts but is markedly lower compared to *APC*$^{+/-}$ midguts that contain *APC*$^{-/-}$clones (*c.f.* Figure 4E). In **(D-F)**, each data point represents one midgut; *n* = 4 midguts per condition. *P* values by unpaired *t*-test. For box-and-whisker plots, the boxes show median, 25$^{th}$ and 75th percentiles, and whiskers are minimum and maximum values.

**Table S1 | Experimental genotypes by figure**

| Figure | Panels | Genotype |
|---|---|---|
| **Figure 1** | Fig. 1B-C | hsflp$^{122}$, UAS-GFP; tubGAL4 / E-cad$^{mTomato}$ ; FRT82B tubGAL80 / TM6B |
| | | hsflp$^{122}$, UAS-GFP; tubGAL4 / E-cad$^{mTomato}$; FRT82B tubGAL80 / FRT82B, APC2$^{G10}$, APC1$^{Q8}$ |
| | Fig. 1F-H | hsflp$^{122}$, UAS-GFP; tubGAL4; FRT82B tubGAL80 / FRT82B |
| | | hsflp$^{122}$, UAS-GFP; tubGAL4 / UAS-E-cad; FRT82B tubGAL80 / FRT82B |
| | | hsflp$^{122}$, UAS-GFP; tubGAL4 / UAS-E-cad$^{dCR4h}$; FRT82B tubGAL80 / FRT82B |
| | | hsflp$^{122}$, UAS-GFP; tubGAL4 / UAS-E-cad$^{\Delta JM}$; FRT82B tubGAL80 / FRT82B |
| | | hsflp$^{122}$, UAS-GFP; tubGAL4 / UAS-p120 RNAi; FRT82B tubGAL80 / FRT82B |
| | | hsflp$^{122}$, UAS-GFP; tubGAL4; FRT82B tubGAL80 / FRT82B, APC2$^{G10}$, APC1$^{Q8}$ |
| | | hsflp$^{122}$, UAS-GFP; tubGAL4 / UAS-E-cad; FRT82B tubGAL80 / FRT82B, APC2$^{G10}$, APC1$^{Q8}$ |
| | | hsflp$^{122}$, UAS-GFP; tubGAL4 / UAS-E-cad$^{dCR4h}$; FRT82B tubGAL80 / FRT82B, APC2$^{G10}$, APC1$^{Q8}$ |
| | | hsflp$^{122}$, UAS-GFP; tubGAL4 / UAS-E-cad$^{\Delta JM}$; FRT82B tubGAL80 / FRT82B, APC2$^{G10}$, APC1$^{Q8}$ |
| | | hsflp$^{122}$, UAS-GFP; tubGAL4 / UAS-p120 RNAi; FRT82B tubGAL80 / FRT82B, APC2$^{G10}$, APC1$^{Q8}$ |
| **Figure 2** | Fig. 2B-D | hsflp$^{122}$; rho-lacZ (rho$^{X81}$), FRT82B, ubi-GFP / FRT82B |
| | | hsflp$^{122}$; rho-lacZ (rho$^{X81}$), FRT82B, ubi-GFP / FRT82B, APC2$^{G10}$, APC1$^{Q8}$ |
| | Fig. 2E-F | hsflp$^{122}$, UAS-GFP; tubGAL4; FRT82B tubGAL80 / FRT82B |
| | | hsflp$^{122}$, UAS-GFP; tubGAL4; FRT82B tubGAL80 / FRT82B, APC2$^{G10}$, APC1$^{Q8}$ |
| | Fig. 2G-I | hsflp$^{122}$, UAS-GFP; tubGAL4; FRT82B tubGAL80 / FRT82B |
| | | hsflp$^{122}$, UAS-GFP; tubGAL4 / UAS-egfr RNAi; FRT82B tubGAL80 / FRT82B |
| | | hsflp$^{122}$, UAS-GFP; tubGAL4 / UAS-rho RNAi; FRT82B tubGAL80 / FRT82B |
| | | hsflp$^{122}$, UAS-GFP; tubGAL4; FRT82B tubGAL80 / FRT82B, APC2$^{G10}$, APC1$^{Q8}$ |
| | | hsflp$^{122}$, UAS-GFP; tubGAL4 / UAS-egfr RNAi; FRT82B tubGAL80 / FRT82B, APC2$^{G10}$, APC1$^{Q8}$ |
| | | hsflp$^{122}$, UAS-GFP; tubGAL4 / UAS-rho RNAi; FRT82B tubGAL80 / FRT82B, APC2$^{G10}$, APC1$^{Q8}$ |
| **Figure 3** | Fig. 3A-B | hsflp$^{122}$; rho-lacZ (rho$^{X81}$), FRT82B, ubi-GFP / FRT82B |
| | | hsflp$^{122}$; rho-lacZ (rho$^{X81}$), FRT82B, ubi-GFP / FRT82B, APC2$^{G10}$, APC1$^{Q8}$ |
| | Fig. 3D-F | hsflp$^{122}$; FRT82B, GS2326, ubi-RFP / FRT82B |
| | | hsflp$^{122}$; UAS-rho RNAi; FRT82B, GS2326, ubi-RFP / FRT82B |
| | | hsflp$^{122}$; UAS-E-cad; FRT82B, GS2326, ubi-RFP / FRT82B |
| | | hsflp$^{122}$; UAS-p120 RNAi; FRT82B, GS2326, ubi-RFP / FRT82B |
| | | hsflp$^{122}$; UAS-egfr RNAi; FRT82B, GS2326, ubi-RFP / FRT82B |
| | | hsflp$^{122}$; FRT82B, GS2326, ubi-RFP / FRT82B, APC2$^{G10}$, APC1$^{Q8}$ |
| | | hsflp$^{122}$; UAS-rho RNAi; FRT82B, GS2326, ubi-RFP / FRT82B, APC2$^{G10}$, APC1$^{Q8}$ |
| | | hsflp$^{122}$; UAS-E-cad; FRT82B, GS2326, ubi-RFP / FRT82B, APC2$^{G10}$, APC1$^{Q8}$ |
| | | hsflp$^{122}$; UAS-p120 RNAi; FRT82B, GS2326, ubi-RFP / FRT82B, APC2$^{G10}$, APC1$^{Q8}$ |
| | | hsflp$^{122}$; UAS-egfr RNAi; FRT82B, GS2326, ubi-RFP / FRT82B, APC2$^{G10}$, APC1$^{Q8}$ |
| **Figure 4** | Fig. 4B-E | hsflp$^{122}$; rho-lacZ (rho$^{X81}$), FRT82B, ubi-GFP / FRT82B, APC2$^{G10}$, APC1$^{Q8}$ |
| | Fig. 4F | hsflp$^{122}$; FRT82B, GS2326, ubi-RFP / FRT82B (reference cDNA) |
| | Fig. 4F-H | hsflp$^{122}$; FRT82B, GSG2326, ubi-RFP / FRT82B, APC2$^{G10}$, APC1$^{Q8}$ |
| | | hsflp$^{122}$ / UAS-bsk$^{DN}$; FRT82B, GSG2326, ubi-RFP / FRT82B, APC2$^{G10}$, APC1$^{Q8}$ |
| **Figure S1** | Fig. S1B | hsflp$^{122}$, UAS-GFP; tubGAL4; FRT82B tubGAL80 / FRT82B, APC2$^{G10}$, APC1$^{Q8}$ |
| | Fig. S1D-E | hsflp$^{122}$, UAS-GFP; tubGAL4 / E-cad$^{mTomato}$ ; FRT82B tubGAL80 / TM6B |
| | | hsflp$^{122}$, UAS-GFP; tubGAL4 / E-cad$^{mTomato}$; FRT82B tubGAL80 / FRT82B, APC2$^{G10}$, APC1$^{Q8}$ |
| | Fig. S1F, H | hsflp$^{122}$; rho-lacZ (rho$^{X81}$), FRT82B, ubi-GFP / FRT82B, APC2$^{G10}$, APC1$^{Q8}$ |
| | Fig. S1G, I | hsflp$^{122}$, UAS-GFP; tubGAL4; FRT82B tubGAL80 / FRT82B, APC2$^{G10}$, APC1$^{Q8}$ |

**Table S1 | Experimental genotypes by figure (continued)**

| Figure | Panels | Genotype |
|---|---|---|
| **Figure S2** | Fig. S2A, D | *hsflp$^{122}$, UAS-GFP; tubGAL4 / UAS-X; FRT82B tubGAL80 / FRT82B, APC2$^{G10}$, APC1$^{Q8}$* |
| | Fig. S2B | *hsflp$^{122}$; FRT82B, ubi-GFP / FRT82B, APC2$^{G10}$, APC1$^{Q8}$* |
| | Fig. S2C, E | *hsflp$^{122}$; UAS-X; FRT82B, GSG2326, ubi-RFP / FRT82B, APC2$^{G10}$, APC1$^{Q8}$* |
| **Figure S3** | Fig. S3A-B | *hsflp$^{122}$; rho-lacZ (rho$^{X81}$), FRT82B, ubi-GFP / FRT82B* |
| | Fig. S3B | *hsflp$^{122}$; rho-lacZ (rho$^{X81}$), FRT82B, ubi-GFP / FRT82B, APC2$^{G10}$, APC1$^{Q8}$* |
| | Fig. S3C-D | *hsflp$^{122}$, UAS-GFP; tubGAL4; FRT82B tubGAL80 / FRT82B, APC2$^{G10}$, APC1$^{Q8}$* |
| | | *hsflp$^{122}$, UAS-GFP; tubGAL4 / UAS-p35; FRT82B tubGAL80 / FRT82B, APC2$^{G10}$, APC1$^{Q8}$* |
| | Fig. S3E-F | *hsflp$^{122}$; FRT82B, GS2326, ubi-RFP / FRT82B, APC2$^{G10}$, APC1$^{Q8}$* |
| | | *hsflp$^{122}$; UAS-p35; FRT82B, GS2326, ubi-RFP / FRT82B, APC2$^{G10}$, APC1$^{Q8}$* |
| | Fig. S3G-I | *hsflp$^{122}$, UAS-GFP; tubGAL4; FRT82B tubGAL80 / FRT82B* |
| | | *hsflp$^{122}$, UAS-GFP / UAS-bsk$^{DN}$; tubGAL4; FRT82B tubGAL80 / FRT82B* |
| | | *hsflp$^{122}$, UAS-GFP; tubGAL4 / UAS-p35; FRT82B tubGAL80 / FRT82B* |
| | | *hsflp$^{122}$, UAS-GFP; tubGAL4; FRT82B tubGAL80 / FRT82B, APC2$^{G10}$, APC1$^{Q8}$* |
| | | *hsflp$^{122}$, UAS- GFP / UAS-bsk$^{DN}$; tubGAL4; FRT82B tubGAL80 / FRT82B, APC2$^{G10}$, APC1$^{Q8}$* |
| | | *hsflp$^{122}$, UAS-GFP; tubGAL4 / UAS-p35; FRT82B tubGAL80 / FRT82B, APC2$^{G10}$, APC1$^{Q8}$* |
| | Fig. S3J-L | *hsflp$^{122}$; FRT82B, GS2326, ubi-RFP / FRT82B* |
| | | *hsflp$^{122}$ / UAS-bsk$^{DN}$; FRT82B, GS2326, ubi-RFP / FRT82B* |
| | | *hsflp$^{122}$; UAS-p35; FRT82B, GS2326, ubi-RFP / FRT82B* |
| | | *hsflp$^{122}$; FRT82B, GS2326, ubi-RFP / FRT82B, APC2$^{G10}$, APC1$^{Q8}$* |
| | | *hsflp$^{122}$ / UAS-bsk$^{DN}$; FRT82B, GS2326, ubi-RFP / FRT82B, APC2$^{G10}$, APC1$^{Q8}$* |
| | | *hsflp$^{122}$; UAS-p35; FRT82B, GS2326, ubi-RFP / FRT82B, APC2$^{G10}$, APC1$^{Q8}$* |
| | Fig. S4A | *hsflp$^{122}$; rho-lacZ (rho$^{X81}$), FRT82B, ubi-GFP / FRT82B* |
| | Fig. S4A-B | *hsflp$^{122}$; rho-lacZ (rho$^{X81}$), FRT82B, ubi-GFP / FRT82B, APC2$^{G10}$, APC1$^{Q8}$* |
| | Fig. S4D-F | *hsflp$^{122}$; rho-lacZ (rho$^{X81}$), FRT82B, ubi-GFP / FRT82B* |
| | | *hsflp$^{122}$; rho-lacZ (rho$^{X81}$), FRT82B, ubi-GFP / FRT82B, APC2$^{G10}$, APC1$^{Q8}$* |

**Table S1. Experiment genotype by figure. Related to STAR Methods.**

*'GS2326'* denotes the GeneSwitch driver, *P[Switch]GSG2326* [S3].

'*UAS-X*' generically denotes a UAS-driven transgene of interest (ex: *UAS-E-cad*).

| Table S2 \| Clone numbers shown per figure panel | | | | | |
|---|---|---|---|---|---|
| | | Control clones | | $APC^{-/-}$ clones | |
| Figure panel | Condition | Total clones | No. of guts | Total clones | No. of guts |
| Figure 1C | Single-layered | - | - | 67 | 4 |
| | Multi-layered | - | - | 38 | 4 |
| Figure 1G | Clones alone | 90 | 3 | 79 | 3 |
| | *E-cad* | 88 | 3 | 79 | 3 |
| | *E-cad*$^{dCR4h}$ | 70 | 3 | 97 | 3 |
| | *E-cad*$^{\Delta JM}$ | 70 | 3 | 80 | 4 |
| | *p120* RNAi | 68 | 3 | 116 | 3 |
| Figure 2C | *rho-lacZ* | 81 | 3 | 47 | 4 |
| Figure 2F | dpERK | 88 | 3 | 74 | 3 |
| Figure 2H | Clones alone | 79 | 3 | 74 | 3 |
| | *egfr* RNAi | 24 | 3 | 53 | 4 |
| | *rho* RNAi | 84 | 3 | 83 | 3 |
| Figure 3E | Clones alone | 75 | 3 | 65 | 4 |
| | *egfr* RNAi | 39 | 4 | 71 | 3 |
| | *E-cad* | 52 | 3 | 72 | 3 |
| | *p120* RNAi | 49 | 3 | 78 | 3 |
| | *rho* RNAi | 76 | 4 | 81 | 3 |
| Figure 4G | Clones alone | - | - | 26 | 3 |
| | *bsk*$^{DN}$ | - | - | 53 | 3 |
| Figure S1G | Single-layered | - | - | 69 | - |
| | Multi-layered | - | - | 31 | - |
| Figure S1H | Single-layered | - | - | 35 | - |
| | Multi-layered | - | - | 12 | - |
| Figure S1I | Single-layered | - | - | 46 | - |
| | Multi-layered | - | - | 28 | - |
| Figure S3D | Clones alone | - | - | 60 | 3 |
| | *p35* | - | - | 53 | 3 |
| Figure S3F | Clones alone | - | - | 29 | 3 |
| | *p35* | - | - | 64 | 3 |
| Figure S3H | Clones alone | 79 | 3 | 40 | 3 |
| | *p35* | 39 | 3 | 53 | 3 |
| | *bsk*$^{DN}$ | 60 | 3 | 48 | 3 |
| Figure S3K | Clones alone | 48 | 3 | 55 | 4 |
| | *p35* | 70 | 3 | 64 | 3 |
| | *bsk*$^{DN}$ | 79 | 3 | 53 | 3 |

**Table S2. Clone numbers shown per figure panel. Related to STAR Methods.**
One of three repetitions is shown for each experiment; numbers of clones per experiment vary between repetitions. For Figure 1C, both early- and late-stage tumors were quantified from the same midguts (n = 4).

| Table S3 \| Experimental values by condition | | | | | |
|---|---|---|---|---|---|
| **Condition** | | **Control clones** | | ***APC**$^{-/-}$* **clones** | |
| **Expression** | **UAS** | **Median clone size** | **% multilayering (mean ± SD)** | **Median clone size** | **% multilayering (mean ± SD)** |
| In clones | No UAS | 8.5 | 0 ± 0 | 37 | 41.4 ± 6.3 |
| | *E-cad* | 3 | 0 ± 0 | 3 | 7.1 ± 3.2 |
| | *E-cad*$^{\Delta JM}$ | 14 | 0 ± 0 | 25 | 39.0 ± 6.4 |
| | *E-cad*$^{dCR4h}$ | 10 | 0 ± 0 | 5 | 7.2 ± 2.8 |
| | *p120* RNAi | 7 | 0 ± 0 | 6 | 6.0 ± 2.2 |
| | *rho* RNAi | 5 | 0 ± 0 | 13.5 | 13.3 ± 2.8 |
| | *egfr* RNAi | 2 | 0 ± 0 | 2 | 0 ± 0 |
| | *p35* | 5 | 0 ± 0 | 20 | 31.1 ± 9.2 |
| | *bsk*$^{DN}$ | 7.5 | 0 ± 0 | 20 | 29.5 ± 6.3 |
| In non-clone cells | No UAS | 9 | 0 ± 0 | 37 | 33.0 ± 4.2 |
| | *E-cad* | 5 | 0 ± 0 | 7 | 9.8 ± 7.4 |
| | *p120* RNAi | 5 | 0 ± 0 | 7 | 11.8 ± 10.2 |
| | *rho* RNAi | 6 | 0 ± 0 | 10 | 15.5 ± 5.8 |
| | *egfr* RNAi | 10 | 0 ± 0 | 42 | 39.8 ± 1.7 |
| | *p35* | 4 | 0 ± 0 | 9.5 | 11.0 ± 7.0 |
| | *bsk*$^{DN}$ | 9 | 0 ± 0 | 12 | 13.0 ± 1.6 |

**Table S3. Experimental values by condition. Related to STAR Methods.**

Values from one of three repetitions is shown for each experiment**.**
